# adequaSSE: Model Adequacy Testing for Trait-Dependent Diversification Models

**DOI:** 10.1101/2023.03.06.531416

**Authors:** Orlando Schwery, Will Freyman, Emma E. Goldberg

## Abstract

The presence of large variation in speciation and extinctin rates across the tree of life has long been hypothesized to be driven by the evolution of traits that affect diversification. To test such hypotheses, phylogenetic biologists have developed a wide class of state-dependent birth-death processes that jointly model trait evolution and the diversification process. However, it has since been shown that these models are sensitive to falsely linking traits to diversification. Here we present a Bayesian approach to test the adequacy of statedependent birth-death models by statistically checking whether they describe the variation observed in the data. Our method generates posterior predictive distributions for a suite of informative test statistics, providing a general framework for testing diversification processes and models of trait evolution. We simulate data sets under different violations of model assumptions and find that our approach successfully detects the inadequacy of the model for them. We further show that the manner in which a model fails to fit the data can reveal insights into the processes of trait evolution and diversification.

## Introduction

To understand why rates of speciation and extinction vary remarkably across the tree of life, evolutionary biologists have developed concepts such as adaptive zones (Simpson 1953), key innovations (Hodges and Arnold 1995), adaptive radiation (Schluter 2000), and species selection (Stanley 1975; Jablonski 2008). The core of these ideas is that a lineage aquiring an advantageous trait in the right ecological context gains a competitive advantage that translates into either higher rates of speciation, lower risk of extinction, or both. Testing these concepts in a macroevolutionary context requires testing the association between diversification rate shifts and trait evolution. Despite enormous progress in the data and methods available for such tests, however, it remains challenging to establish a robust case for trait-dependent diversification.

The most powerful methods would seem to be those that explicitly model the processes of speciation, extinction, and trait evolution, such as a multi-type branching process (Maddison et al. 2007, and many variants thereof). However, such models can associate diversification effects with traits that do not warrant them (Rabosky and Goldberg 2015) when the models do not allow that other processes could explain diversification rate heterogeneity that is in the data (Beaulieu and O'Meara 2016). One response is to create more complicated models that do include other processes affecting diversification (Beaulieu and O'Meara 2016; Caetano et al. 2018; Zenil-Ferguson et al. 2019). Such models become increasingly difficult to fit to data, though, and it is never known when they include ‘enough.’

An alternative, which we pursue here, is to test whether a given model is statistically adequate for the data at hand, rather than merely testing whether it fits better than alternative models. Adequacy testing has been adopted in other areas of phylogenetics, such as alignment of nucleotide and amino acid sequences (Huelsenbeck et al. 2001; Nielsen and Huelsenbeck 2001), models of sequence evolution (Nielsen 2002; Bollback 2002; Brown and ElDabaje 2008; Lewis et al. 2014), continuous trait evolution (Slater and Pennell 2014; Pennell et al. 2015), discrete trait evolution (Huelsenbeck et al. 2003; Blackmon and Demuth 2014), tree branch lengths and topologies under phylogenetic inference (Brown 2014) and the birth-death process (Schwery and O'Meara 2020), the multispecies coalescent (Reid et al. 2014), and phylodynamics (Duchene et al. 2018). For the topic of trait-dependent lineage diversification, Bromham et al. (2016) and Ng and Smith (2018) performed adequacy assessments targeted at dead-end traits, but no general framework exists. This is what we develop.

Here, we perform posterior predictive checking of model adequacy for BiSSE models. Inadequacy of the model is indicated when test statistics calculated from the observed data are outliers compared to the same test statistics calculated from data simulated under the fitted model (Rubin 1984; Gelman et al. 1996). We employ a very large set of statistics that summarize different aspects of tree shape and trait distributions, and we assess how well these perform for adequacy testing. We find that suites of test statistics can highlight when the fitted model does not match the processes under which the data were generated. Furthermore, we show that different magnitudes of inadequacy can be revealed, and that different processes causing inadequacies can be distinguished. We close by highlighting the distinction between two different types of adequacy: no model will ever contain all the processes under which real-world data are generated (all models are inadequate for the data), but a model may still provide valid inferences about trait-dependent lineage diversification (some models are adequate for the question).

## Methods

### General approach

The core of our approach is to determine the adequacy of a candidate model for a given observed dataset by assessing whether the model is able to reproduce the key characteristics of the data, i.e., whether model-generated data ‘looks like’ the observed data. We achieve this using posterior predictive simulations (PPS; Fig. 1, Gelman et al. (1996)). First, the model is fit to the data to estimate the model parameters (speciation, extinction, and state transition rates, in case of BiSSE). This primary inference step is done in a Bayesian framework, yielding posterior distributions of the parameters which include their uncertainties and correlations. Second, a set of model parameter values is drawn from the posterior distribution, and those values are used to simulate data (a tree and tip character states, in our case). Repeating this for many draws of parameter values generates a large set of simulated data. Third, on the observed dataset and each simulated dataset, a variety of summary statistics are calculated. These summary statistics are intended to capture the relevant dimensions of variation across the data. Fourth, for each summary statistic, the value from the observed data is compared against the distribution across the simulated data. Posterior predictive *p*-values describe whether the observed data is an outlier relative to the data simulated under the model. Finally, the *p*-values of all summary statistics are interpreted to conclude whether the employed model is adequate for the data at hand. Summary statistics that have particularly low *p*-values (i.e., for which the observed data is particularly far from what the model produced) may hint at processes involved in generating the observed data but not present in the fitted model.

**Figure 1:**
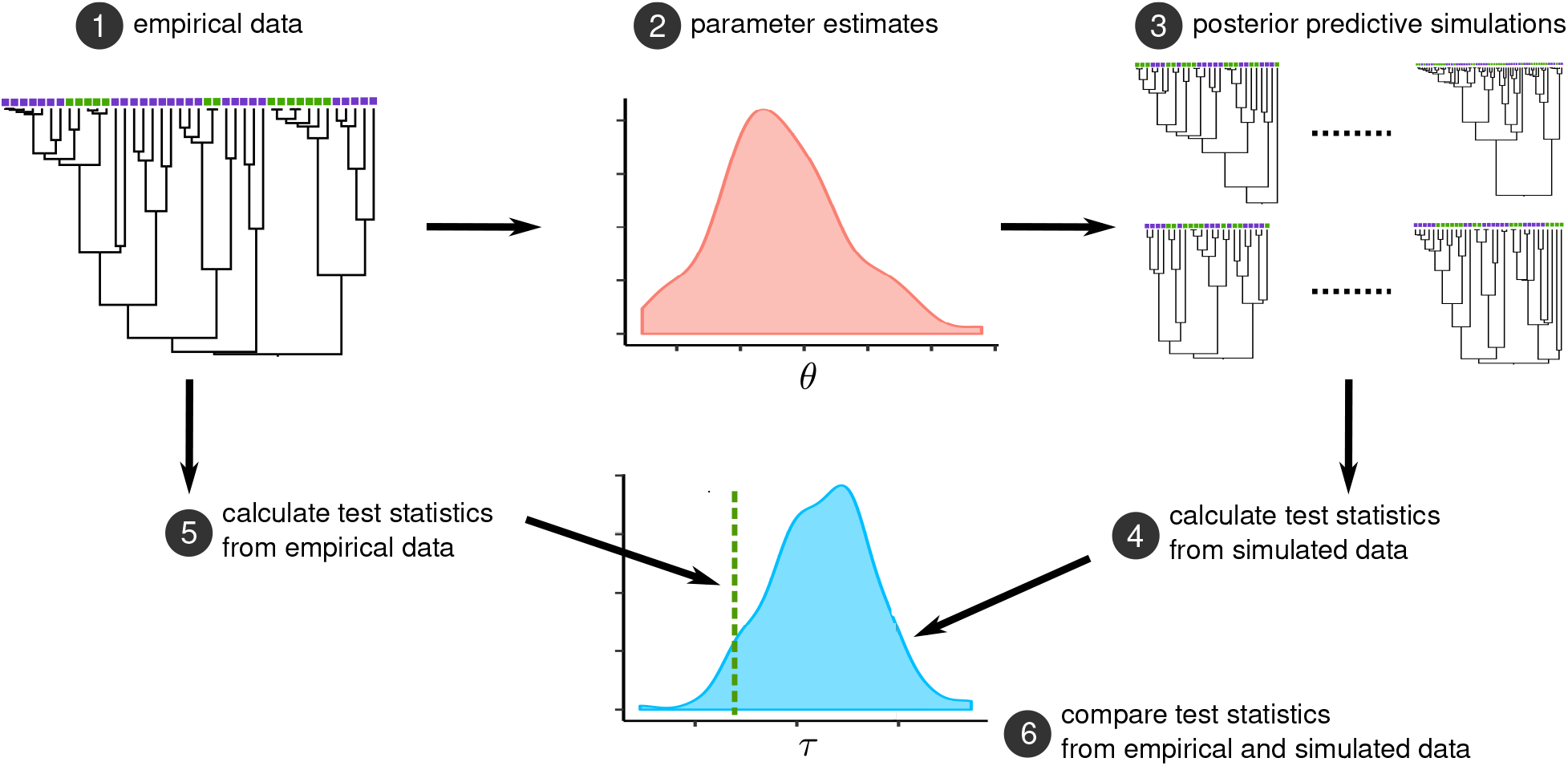
Posterior predictive checks for state-dependent diversification models. (1) Start with observed data, here a tree and tip states. (2) Fit the inference model to obtain the posterior distribution of its parameters. (3) Simulate many data sets, using the inference model as the generating model, with parameter values drawn from the posterior. (4) Calculate each test statistic for all the simulated datasets, producing its posterior predictive distribution. (5) Calculate the same test statistics from the observed data. (6) Compare each observed test statistic value against its simulated distribution. If the test statistic calculated from the observed data is an outlier compared to the posterior predictive distribution, than the model is considered inadequate. This is quantified by calculating posterior predictive *p*-values for each test statistic.

Several challenges arise when applying this standard PPS approach to trait-dependent diversification, in which the generating model includes both evolution of the trait and the growth of the tree itself. The distribution of data simulated under a birth-death process can be extremely broad, so we impose some constraints by conditioning the simulations on various specific features of the data. We must also select summary statistics that capture the distribution of traits, the shape of the tree, and their interaction. And in interpreting the findings across all summary statistics, we must consider what exactly we want to learn from a model adequacy test. No inference model will truly be adequate to explain all the intricacies of any empirical dataset, but we can instead aim to determine whether an inference model is adequate to test the association of a trait with lineage diversification rates. Each of these challenges is discussed further below.

### Inference models

Our explorations here include two commonly-used models as the inference model. The simpler is a constant-rate birth-death model (Nee et al. 1994) for the tree, on which evolves a binary trait that is not associated with the speciation or extinction rate (Pagel 1999). We call this model CID-1 because it is a character-independent diversification model with one trait (Beaulieu and O'Meara 2016). The other is the BiSSE model, in which speciation and extinction rates depend on the state of an evolving binary trait (Maddison et al. 2007). More complex inference models can be used to assess whether trait-dependent diversification reported by BiSSE is robust to the action of additional processes (e.g., CID-2 and HiSSE; Beaulieu and O'Meara 2016). Extending our investigations to such models presents a substantial computational burden but no additional conceptual complications. We do not pursue it here, however, because our central goal is to develop a procedure—using PPS rather than statistical model comparisons—to determine whether BiSSE is an adequate model for testing the association of a trait with diversification rate. We consider a much wider array of generating models, described further below.

### Conditioning posterior predictive simulations on the data

Due to the stochasticity inherent in the birth-death process, one set of diversification rates will yield a wide range of phylogenies which differ along many dimensions such as tree age, number of tips, or tree balance. Some summary statistics may consequently take on posterior distributions that are so broad as to be powerless for assessing adequacy. We therefore constrain the variation in our simulations by conditioning on certain aspects of the observed data. Imposing different constraints produces different distributions of some summary statistics, potentially providing different sensitivities. Our simulations each constrain one of: the start time of the tree (the time duration of the forward-time birth-death process), the number of surviving tips (the stopping condition of the forward-time birth-death process), the number of tips in each trait state (simulated backward in time, Bromham et al. 2016 for BiSSE), or the tree itself and the tip states (via stochastic mapping, Freyman and Höhna 2019 for BiSSE). Technical details are provided in Appendix A.

For example, if we condition our simulations on the age of our tree, the number of surviving lineages could be an informative summary statistic. Alternatively, if we condition on the number of taxa, the number of surviving lineages will be an entirely uninformative summary statistic, but the tree’s age could be informative. Either of those conditionings may also yield useful information from summary statistics that incorporate the distribution of trait states across the trees. Instead conditioning on the number of lineages in each trait state still fixes the number of extant tips in the tree and also reduces variation in their states. This obviously renders the state frequency at the tips an uninformative summary statistic, but it may make more useful the summary statistics that combine trait with branch length. Finally, we can reduce the available space for the simulations even more by conditioning on the observed tree and its tip states. Here, all summary statistics concerned with only topology or branch lengths are also uninformative. Summary statistics using the number of inferred trait origins, however, may be given more power to reveal differences between models. These examples illustrate how we can regard the same summary statistic under a different conditioning as if it were a different summary statistic altogether.

### Summary statistics

For models of diversification and trait evolution, we intuitively expect that the tree topology, its branch lengths, and the distribution of trait states are all important. There is no fixed procedure for distilling these attributes into specific computable summary statistics, however, so experimentation is required. Our strategy is to begin with a large panel of summary statistics and explore their usefulness for our questions.

Table 1 summarizes the types of summary statistics and the variants we employ. Some statistics are direct measures of the overall tree such as tree length, number of surviving lineages, frequency of trait states at the tips, or simple characterizations of the distributions of branch lengths and node ages. Others involve trait reconstructions, such as the total branch lengths in a trait state according to stochastic mapping or parsimony reconstruction, or the parsimony score. Still others combine tree shape and tip states in ways derived from diversification studies and community phylogenetics.

**Table 1:**
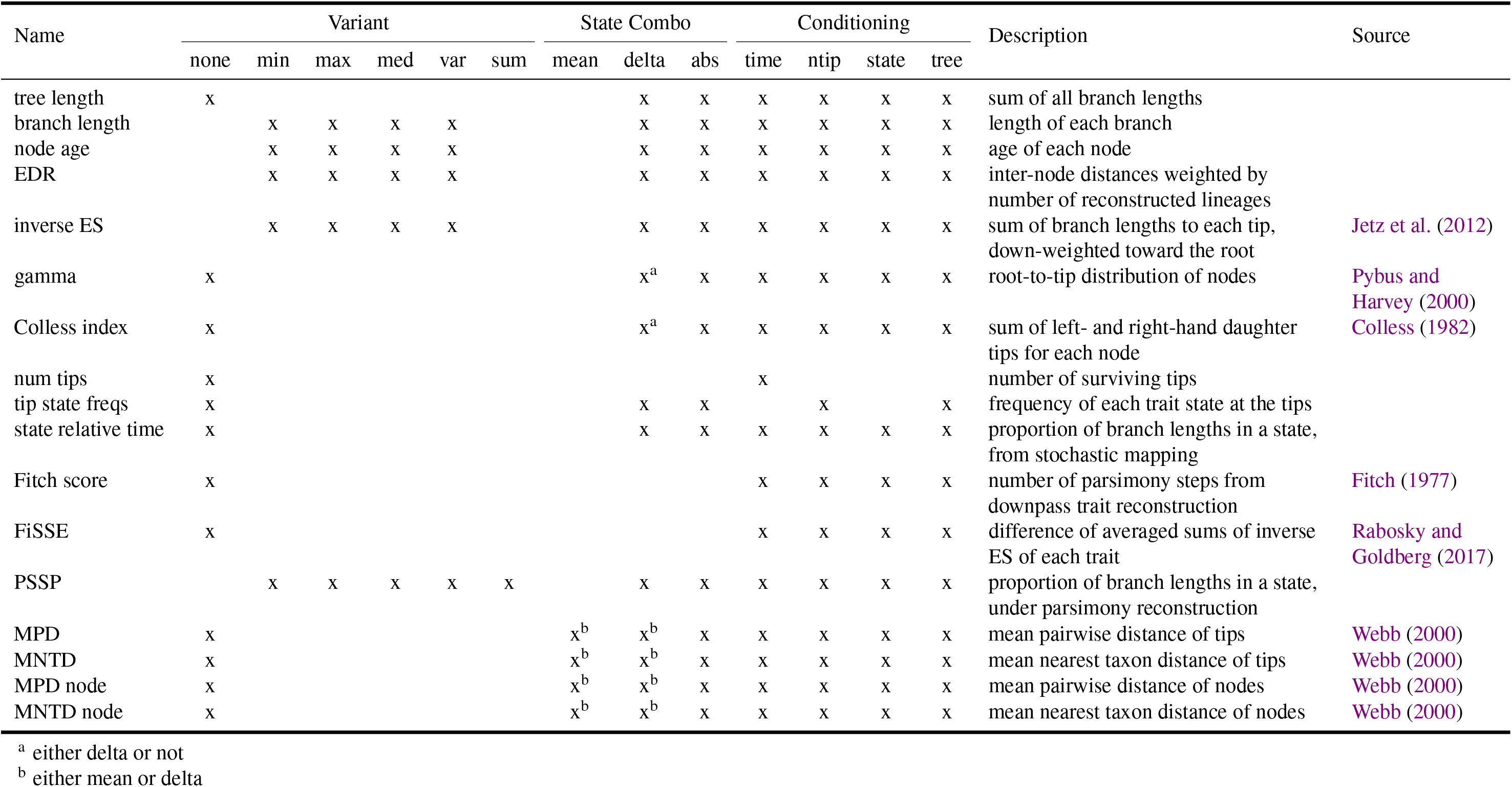
Summary statistics used for model adequacy testing. Each statistic was either calculated as a single value (no variants) or as a set of values which were then summarized as minimum, maximum, median, variance, or sum. For statistics computed on trees pruned to contain tips of just one state, values for the two states were combined as either the mean across the two states, the difference, or the absolute value of the difference. Posterior simulations were conditioned on one of four criteria: time of crown age, number of extant tips, number of taxa in each state, or tree fixed to input phylogeny (only trait is simulated); not all conditionings are logically possible for all statistics.

Many of these summary statistics are inherently only a single value for a given tree. For others that are derived from a set of values for one tree (e.g., the ages of all nodes on the tree), we use the minimal, maximal, median, mean, or variance of the values. We also incorporate trait information by computing additional variants for many summary statistics through pruning the tree to only include the tips of either of the binary trait states, calculating the summary statistic in question for each pruned tree, and recording the difference (‘delta’) and absolute difference (‘absdelta’) between them. Additionally, some summary statistics make sense only under some conditionings, as discussed above. The complete set of 269 statistics we considered is given in Table S1.

### Posterior predictive p-values

The PPS procedure calculates the values of a summary statistic on simulated trees to compose an empirical density function under the inference model. The tail area of that distribution, measured from the statistic’s value on the observed tree, is the posterior predictive *p*-value. If this value is small, the model is presumably not a good fit to the data. The logic is similar to a frequentist *p*-value, but posterior predictive *p*-values are not calibrated to reflect the probability of the data under the model. In particular, distributions of posterior predictive *p*-values tend not to be flat, but rather to be concentrated away from 0 and 1 due to uncertainty in the posterior (Meng 1994; Gelman 2013). Consequently, if we declare that a model is inadequate for our data when the posterior predictive *p*-value is less than, say, 0.05, we will be conservative in reporting inadequacy—likely fewer than 5% of datasets simulated under the generating model would be declared inadequate. It would be possible to use simulations to determine the tail probability that corresponds to a false positive rate of 0.05 for a summary statistic. This would need to be re-done for each observed dataset, and it may or may not be desireable (Gelman 2013). In this study, we are working with thousands of ‘empirical’ datasets—replicates under many scenarios, described below—so calibrating the false positive rate for each is impractical. Therefore, we proceed with the convention that a posterior predictive *p*-value of less than 0.05 indicates a poor fit of the model to the data, knowing that this probably reduces the sensitivity of our adequacy test.

### Simulations and tests

In order to test the usefulness and reliability of our summary statistics in determining model adequacy, we simulated test data under a variety of models and scenarios. Our primary focus in this paper is distinguishing the presence of trait-dependent diversification from its absence, and the sensitivity of the test to different kinds and magnitudes of violations of the assumptions of the basic BiSSE model.

Initially, we wanted to confirm whether a model is reliably deemed to be adequate for data that were generated under that same model. We therefore simulated data under our two inference models, representing trait evolution without diversification rate heterogeneity (CID-1) and trait-dependent diversification (BiSSE). We excluded from subsequent analyses any summary statistics that incorrectly signaled model inadequacy on more than 5% of the datasets from the generating model when it was the same as the inference model.

Primarily, we wanted to test how well our method performs on cases where the assumptions of the BiSSE model were violated in different ways and to different extent. Empirical phylogenies are the result of numerous processes, all complicating the picture of trait-dependence by adding various kinds of background rate heterogeneity that is capable of misleading BiSSE. It would thus be very valuable to know whether we are able to detect the presence of such processes rendering the BiSSE model inadequate. And it would be even more valuable if we could leverage our summary statistics to distinguish different kinds of violations of BiSSE’s assumptions. To this end, we simulated many datasets under diversification scenarios that either add an additional process, add more complexity to the trait-dependent process, or involve trait-independent rate heterogeneity which may falsely suggest trait-dependence.

First, we made use of trees simulated by Rabosky (2014) under a complex trait-independent model where lineages transition between up to four density-dependent rate regimes (DD-k4). On these trees we simulated the evolution of a diversification-neutral trait. These datasets are of particular interest because they have been demonstrated to be misleading BiSSE to falsely infer trait-dependence when there is clearly none (Rabosky and Goldberg 2017).

Second, we simulated data under a time-dependent BiSSE model (tdBiSSE), in which diversification rates differ between trait states, but the effect varies over time. These scenarios violate BiSSE’s assumption of rates being constant over time, but trait-dependent diversification is still the appropriate conclusion. We considered cases where the rates decrease with time towards the present, either linearly or exponentially (Fig. S1). Rather than encouraging false inferences of trait-dependence, these latter sets of simulations are meant to test our approach’s behaviour in a scenario where there is indeed trait-dependence, but not only.

Comparing results between the DD-k4 and tdBiSSE datasets, we sought to identify any summary statistics that reliably signalled model inadequacy across a variety of processes. We also sought to identify other summary statistics that signalled only one of the types of inadequacy, and thus could be useful for inferring what additional processes—beyond trait-dependent diversification—contributed to a real dataset. Finally, we tested whether our approach is sensitive to quantitative differences in the extent of model violations, by simulating each scenario with a variety of rate magnitudes.

In total, we used 16 different simulation scenarios or generating models (Table 2). For each, we simulated 100 trees of 500 extant taxa. The tdBiSSE datasets were generated using the R package castor (Louca and Doebeli 2018), and the others were generated in RevBayes (Höhna et al. 2014, 2016).

**Table 2:** Parameter values for the simulation scenarios. For each scenario, 100 outcomes were simulated. For DD-k4, traits were simulated on existing trees (100 samples closest to 500 tips each were chosen). For tdBiSSE, the speciation and extinction rates shown represent the starting values from which the rates decayed linearly or exponentially over time (Fig. S1).

| Model | diversification rate | transition rate | $\lambda_0$ | $\lambda_1$ | $\mu_0$ | $\mu_1$ | $q$ | ntips |
| --- | --- | --- | --- | --- | --- | --- | --- | --- |
| CID-1 | moderate | low | 1.00 | 1.00 | 0.30 | 0.30 | 0.10 | 500 |
| CID-1 | moderate | high | 1.00 | 1.00 | 0.30 | 0.30 | 0.90 | 500 |
| CID-1 | high | low | 10.00 | 10.00 | 0.30 | 0.30 | 0.10 | 500 |
| CID-1 | high | high | 10.00 | 10.00 | 0.30 | 0.30 | 0.90 | 500 |
| BiSSE | moderate | low | 1.00 | 2.00 | 0.30 | 0.30 | 0.10 | 500 |
| BiSSE | moderate | high | 1.00 | 2.00 | 0.30 | 0.30 | 0.90 | 500 |
| BiSSE | high | low | 1.00 | 10.00 | 0.30 | 0.30 | 0.10 | 500 |
| BiSSE | high | high | 1.00 | 10.00 | 0.30 | 0.30 | 0.90 | 500 |
| DD-k4 |  | low |  |  |  |  | 0.10 | 498 [181–913] |
| DD-k4 |  | high |  |  |  |  | 0.90 | 498 [181–913] |
| tdBiSSE | moderate, linear | low | 0.50 | 1.50 | 0.10 | 0.10 | 0.10 | 500 |
| tdBiSSE | moderate, linear | high | 0.50 | 1.50 | 0.10 | 0.10 | 0.90 | 500 |
| tdBiSSE | high, linear | high | 1.00 | 10.00 | 0.10 | 0.10 | 0.90 | 500 |
| tdBiSSE | moderate, exponential | low | 0.50 | 1.50 | 0.00 | 0.01 | 0.10 | 500 |
| tdBiSSE | high, exponential | low | 1.00 | 10.00 | 0.00 | 0.01 | 0.10 | 500 |
| tdBiSSE | high, exponential | high | 1.00 | 10.00 | 0.00 | 0.01 | 0.90 | 500 |

## Results

### Selecting robust summary statistics

We expected that fitting the CID-1 model to trees generated under the CID-1 scenarios should not yield indications of model inadequacy, and similarly for fitting the BiSSE model to trees generated under the BiSSE scenarios (which correctly inferred trait-dependence for all trees in all scenarios except 7 trees under moderate diversification and high transition rates). Of the 267 summary statistics that we considered, 86 met these expectations (Table S1). They represent the overlap of 124 summary statistics which were consistently accurate for CID-1 and 177 which consistently were so for BiSSE. For many summary statistics, however (68 for CID-1, 54 for BiSSE, 64 combined), inadequacy of the correct model was only flagged for 5–10% of the simulated trees of one single scenario. So given that the underlying 0.05 cutoff for the predictive *p*-values is likely to be overly strict (Gelman 2013; Meng 1994), it is well possible that our selection of robust summary statistics is rather conservative as well.

### Identifying an inadequate model

We next tested whether the remaining summary statistics can identify data that were not generated under the BiSSE process. It has been shown that the BiSSE model can yield incorrect findings of state-dependent diversification on trees with substantial diversification rate heterogeneity and a rapidly evolving trait that does not affect diversification (Rabosky and Goldberg 2017). On these same trees with a relatively low rate of trait evolution, we found that two summary statistics flagged the BiSSE model as inadequate ~85% of the time, and three more at least 50% of the time (Fig. 2). Increasing the rate of trait evolution, an additional 12 summary statistics flagged inadequacy 70–90% of the time and six more at least 50% of the time. Thus, for data that could otherwise yield an inference of state-dependent diversification with BiSSE (which was indeed the case for 57% and 92% in the low and high transition rate scenarios respectively), model inadequacy can be identified and the inference error can be avoided.

**Figure 2:**
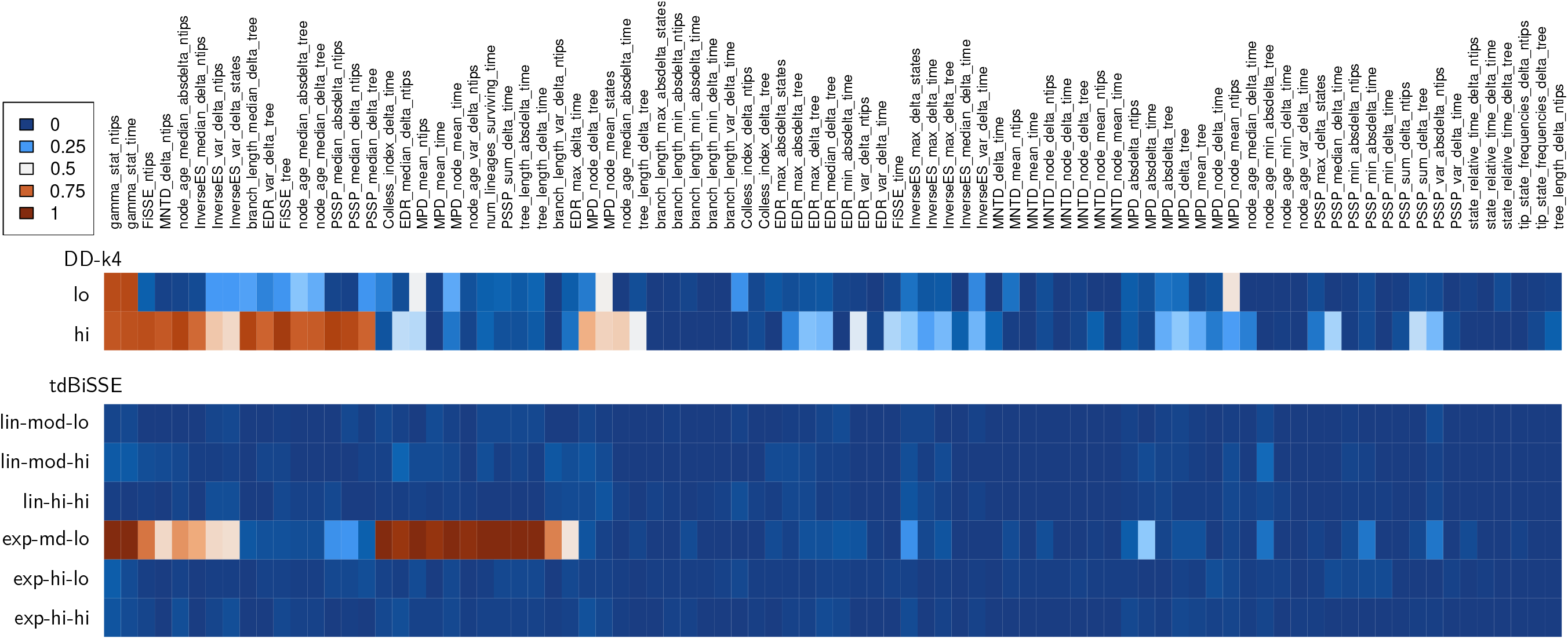
Summary statistic significance for different generating scenarios. For each scenario (DD-k4 is diversification rate heterogeneity, tdBiSSE is time-dependence [Fig. S1]), the proportion of trees is shown for which the summary statistic flagged as significant.

The two most consistent summary statistics to highlight inadequacy for this model were the *γ* statistic conditioned on time or number of tips. This summary statistic captures differences in the distribution of internal nodes across time, independent of the trait. The heterogeneous diversification process that generated these trees is therefore leaving a clear signal in the node ages across the tree. When transition rates are higher, we additionally get alerted to inadequacy by the FiSSE statistic (conditioned on number of tips and the tree), median PSSP between states (conditioned on number of tips, to a lesser degree on the tree), and median differences of branch length and node age between states (conditioned on the tree and number of tips, respectively, node ages to a lesser degree also on the tree). Other statistics with moderate chances of finding adequacy are MPD of nodes and tips, inverse ES, MNTD, and EDR.

### Inadequacy for the data versus inadequacy for the question

No real-world data will be generated under a model as simple as BiSSE, but we still wish to identify traits that do influence diversification. We thus considered scenarios that do include state-dependent diversification, but that also include additional processes not in the simplest BiSSE model. We focused on time-dependent BiSSE processes (tdBiSSE), in which the magnitude of state-dependent diversification varies over time (Fig. S1).

We found that for most of the tdBiSSE scenarios fit with the BiSSE model, inadequacy was not flagged by the summary statistics, with the highest rate at which a statistic indicates inadequacy being 15% (Fig. 2). This is not what we expected, because the inference model is simpler than the generating model and hence cannot correctly capture the processes that generated the data. We suspect that inadequacy in future tests could emerge in two ways: by designing summary statistics that are more sensitive to time-dependent rates, and by designing scenarios that amplify the time-dependent signal.

Indeeed, for the one tdBiSSE scenario that exhibited clear inadequacy of the BiSSE model, the difference in diversification rates was greater over the timespan of the trees (Fig. S1). For this scenario, nine summary statistics correctly signalled inadequacy for every simulated tree. Three further ones signalled more than 90% of the time, and eight between 50–70%. The most reliable summary statistics included Colless between states, tree length between states, the γ statistic, number of surviving lineages, and variants of PSSP and MPD (notably often when conditioned on time).

Our approach thus correctly identifies that the inference model is inadequate for the data. We note, however, that for the research question of whether the trait is associated with diversification, BiSSE is actually an adequate model in these scenarios: it correctly infers significant differences in net diversification rate between trait states for all trees in all tdBiSSE scenarios.

### Distinguishing causes of inadequacy

Beyond simply identifying a model as inadequate, we hoped that the identities of the flagging summary statistics could indicate which processes are present in the data but missing from the model. Some summary statistics flagged inadequacy in both the DD-k4 and tdBiSSE scenarios: the two *γ* statistics (conditioned on number of tips or time) provided a strong signal for both DD-k4 scenarios and were among the most reliable statistics for the tdBiSSE scenario. This suggests that node distribution across time is a general signal of BiSSE inadequacy, not one that can distinguish different causes. Particularly instances of inverse ES (variation and median), but also FiSSE score, MPD, and MNTD (all conditioned on number of tips), and median node age difference between states (conditioned on number of tips) were shared between DD-k4 and tdBiSSE as well. In contrast, other summary statistics flag inadequacy of only one generating process or the other: MPD of nodes, median node age, and median PSSP are more commonly found for DD-k4, while Colless, EDR, surviving lineages, and tree length are more commonly found for tdBiSSE. This division is not strict, and it would need to be investigated across more scenarios, but it does suggest that different summary statistics may have the power to reveal different processes causing model inadequacy.

## Discussion

In this study, we used posterior predictive simulations and a large set of summary statistics to test the adequacy of trait-dependent diversification models for a variety of simulated datasets. We found that our approach can correctly detect the inadequacy of a simple BiSSE model when the generating model includes diversification rate heterogeneity that is independent of trait evolution, and when rates of trait-dependent diversification change over time. We also showed that our approach need not be binary ‘adequate or not,’ but can distinguish varying severity of violations of the inference model’s assumptions.

### Technical improvements

Compared with a parametric bootstrapping approach (Bromham et al. 2016; Ng and Smith 2018), using a posterior predictive framework makes model adequacy tests more conservative—less likely to report inadequacy—because the test incorporates uncertainty in the estimated parameter values. In addition to being Bayesian, our implementation in RevBayes is modular, and in fact some of the components we employed have previously been used for posterior predictive checks on substitution models (Höhna et al. 2018). We further used options that allow for simulating data conditioned on the real data in different ways, effectively generating additional panels of summary statistics. And our workflow can be modified to use other trait-dependent diversification models available in RevBayes, such as HiSSE.

Our results show that the reliable summary statistics differ among models, implying that different processes affect different aspects of the tree (as one might expect). This suggests the need for ‘calibration’ simulations under each inference model (i.e., simulate, infer, and run PPS all under the same model, as we did above), in order to identify which sets of statistics are reliable and meaningful for each model. It remains to be seen whether this extends to paramter space, i.e., whether the usefulness of a summary statistic varies with combinations of model parameter values. If this were to be the case, it might mean that such calibration simulations should be tailored to each specific dataset. Alternatively, a study systematically exploring the performance of summary statistics throughout parameter space for different models may allow subsequent users to inform their analyses, akin to a benchmark database. Such calibration simulations could then also be used as a starting point for determining an α-level of appropriate stringency for each summary statistic.

Of the many summary statistics we considered, relatively few were effective at identifying model inadequacies. Another avenue for improvement is therefore developing additional summary statistics specifically for trait-dependent diversification models. To detect the signature of multiple diversification rate categories in a tree, one might consider changes in node density across the tree (i.e., the autocorrelation of the length of closely related branches), or variation in local γ or Colless values for subtrees. To corroborate the signature of trait dependence in the presence of other processes, the key is to focus on statistics that indicate the validity of the trait-diversification rate association, rather than focusing on the discrepancies from BiSSE’s assumptions. These should incorporate both the tree shape and trait distribution, such as summarizing across the numbers of tips in each state descended from each node, mean clade size per trait (Ng and Smith 2018), and perhaps simple net diversification estimates (Magallon and Sanderson 2001).

### The role of adequacy tests

Adequacy testing can fit into the comparative methods workflow in (at least) two ways. A user could first assess the adequacy of each candidate model, and then compare the remaining adequate models (e.g., via Bayes factors, which are also available in RevBayes) to identify the best-fit model or compute model weights. Alternatively, the best-fit model could first be determined, with its results trusted only if it also proves to be adequate. This could help alleviate errors due to ‘accepting’ a too-simple model solely by rejecting an even-simpler one (Beaulieu and O'Meara 2016). Either way, as usual, the candidate model set should exclude models that are so complex as not to clearly represent a hypothesis of interest. If no model is identified as adequate, however, presumably the models considered should be expanded to include additional processes.

Ideally, the summary statistics that report inadequacy would also indicate the processes that left an imprint on the data but were not included in the inference model. Our results show a hint of this, with a subset of summary statistics flagging inadequacy for one set of generating processes versus another: roughly speaking, the inadequate summary statistics for data generated with neutral rate hetergeneity emphasized the trait distribution and tree balance, while those for data generated with time-dependence more emphasized node ages. Further investigation is needed, however, to establish how consistent such signals can be, and perhaps to design summary statistics intentionally to distinguish specific processes. In the future, model adequacy tests on empirical data might identify processes important in nature but not available in existing methods, thus spurring new model development.

One critical caveat is that failure to be demonstrated inadequate does not prove that a model is adequate, let alone correct. No macroevolutionary model will ever truly capture the diverse biological and geological processes that generated any real clade. Thus, one might argue that a valid test of model adequacy should conclude that any model is inadequate for any dataset. Such a test does not seem very helpful, however. Instead, there is some art to designing a useful adequacy test, such that it is stringent enough to be informative but not so stringent as to be overwhelming negative.

Convincingly testing whether trait-dependent diversification has shaped any real dataset remains a challenge. We know not to reach this conclusion simply because BiSSE fits better than a trivial null model (Beaulieu and O'Meara 2016), but how much complexity must be added to each model to make a rigorous case? We would hope that adequacy testing would be helpful here, but it is not a silver bullet. For example, if HiSSE fits the data better than CID-4, and HiSSE fits adequately but CID-4 does not, there is still no guarantee that trait-dependent diversification is present because other confounding processes may have been missed. Or, if BiSSE fits the data better than CID-1 but neither is adequate, trait-dependent diversification could still be acting along with other processes. What we really need, then, is not so much a test of whether an inference model adequately explains the data, but whether it is adequate to provide a reliable answer to the question at hand (Tiao and Xu 1993; Gelman et al. 1996). We believe this can be achieved through further refinement of the posterior predictive framework used here. The key is to design the interpretation of the summary statistics so as to distinguish not simply whether the inference model is identical to the generating model, but to distinguish a variety of models with trait-dependent diversification (along with other processses) from a variety of models without it.

## Acknowledgements

We thank Tanjona Ramiadantsoa and Heath Blackmon for their help with compiling and implementing summary statistics in the early days of this project. We also thank Jeremy Brown, April Wright, Josef Uyeda, Ethan Romero-Severson, Thomas Leitner, Nick Hengartner, Luke Harmon, Rosana Zenil-Ferguson, and Matt Pennell for helpful discussions at various stages of the process. This project was supported by NSF grant DEB-1655478/1940868 to EEG.

## Appendix A: Posterior Predictive Simulation

The posterior distribution of an assumed model 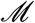 is given by Bayes’ rule,

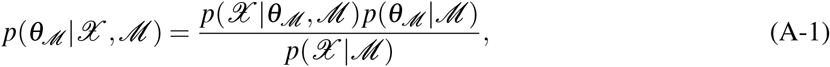

where 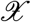 is the observed data, 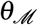 are the parameters of the model, 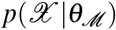 is the likelihood of the model, 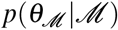 is the joint prior distribution of the model parameters, and 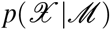 is the marginal likelihood. Typically samples from the posterior distribution 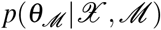 are drawn using Markov chain Monte Carlo (MCMC) algorithms. We are interested in generating the posterior predictive distribution 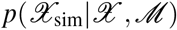, which is the probability of a new data set 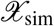 given the observed data set 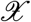 and the model 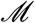. This is defined by

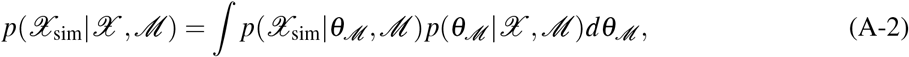

where 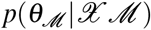 is the posterior distribution estimated using MCMC. Samples from 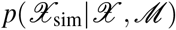 can be generated by simulating new data sets 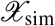 under the model with parameter values 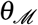 sampled from the posterior distribution.

### Conditioning on aspects of the data

In the case of SSE models, 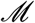 is a joint model of both the birth-death process that generates the phylogeny itself and a character evolving along the branches of the phylogeny. Our observed data 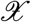 may consist of either the tree *ψ*_obs_, the character data 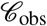, or both. To test different aspects of the data generating process, different posterior predictive distributions may be approximated using Monte Carlo simulations that condition on different parts of the observed data.

The posterior predictive distribution 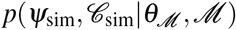, consists of simulated trees *ψ*_sim_ and simulated character data sets 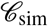 conditioned on the posterior parameter values 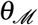 and the model 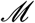, but on no additional features of the data. Due to the stochastic nature of birth-death processes, trees from this distribution will have an extremely large variance and will likely not be informative to distinguish among different SSE processes.

We might instead calculate the posterior predictive distribution 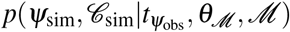 which additionally conditions on the root age *t*_ψobs_ and will therefore generate a narrower distribution of trees. The procedure for PPS conditioned on root age is described in Algorithm 1.

#### Algorithm 1 Forward-time simulation of the state-dependent birth-death process conditioned on the age of the root node. 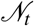 is the vector of the number of lineages in each state at time *t*. The root age is at time *t_r_* > ^0^ and the present is at time *t* = 0.

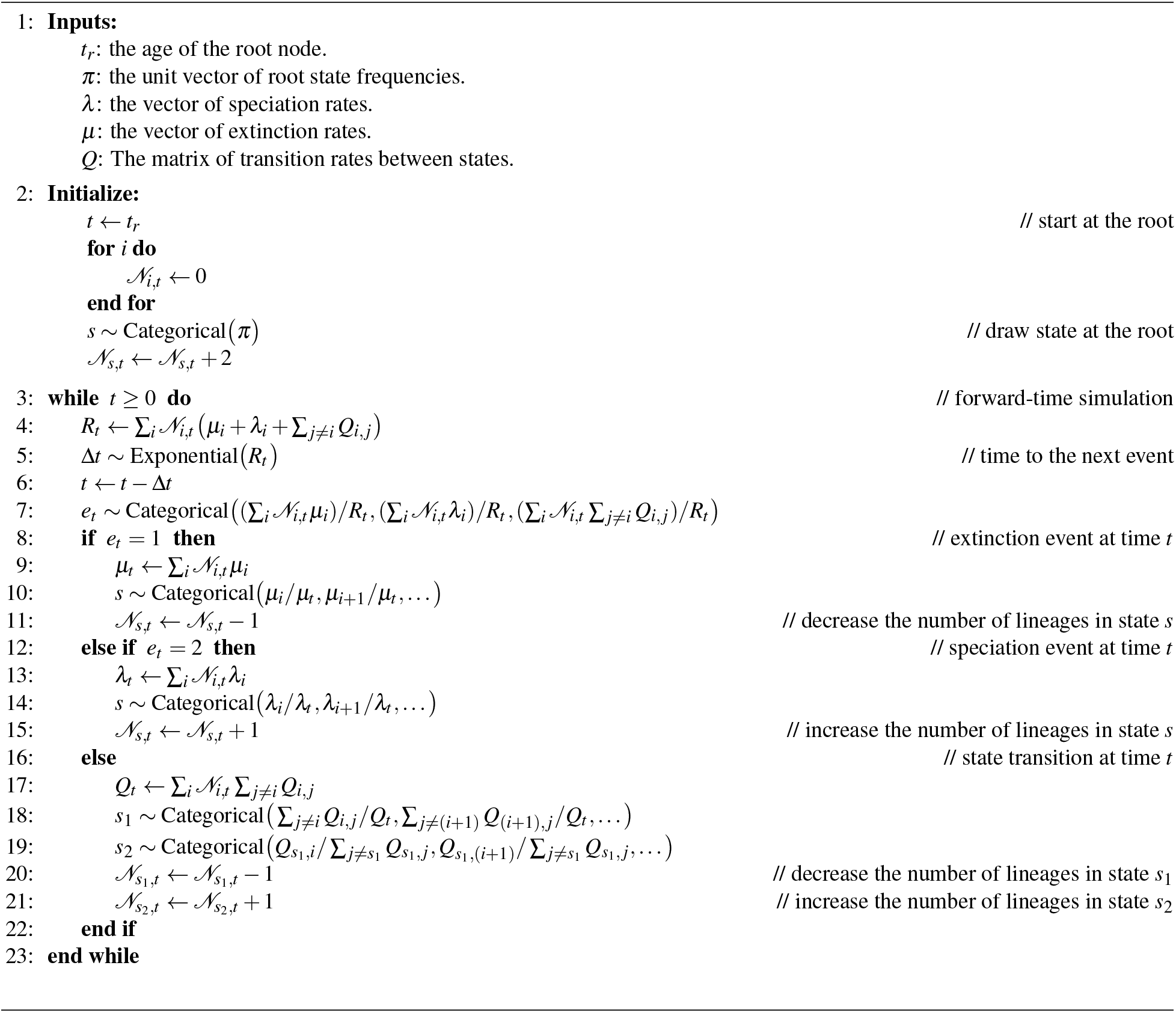

Alternatively, we could calculate 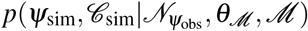 which conditions on the number of tips of the observed tree, 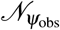. This will also generate a narrower distribution of trees, but in a different manner than conditioning on the root age. The procedure for PPS conditioned on number of tips is described in Algorithm 2.

#### Algorithm 2 Forward-time simulation of the state-dependent birth-death process conditioned on the number of lineages surviving at the present. 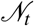 is the vector of the number of lineages in each state at time *t*. The root age is at time *t* = 0 and the present is at some time *t* < 0.

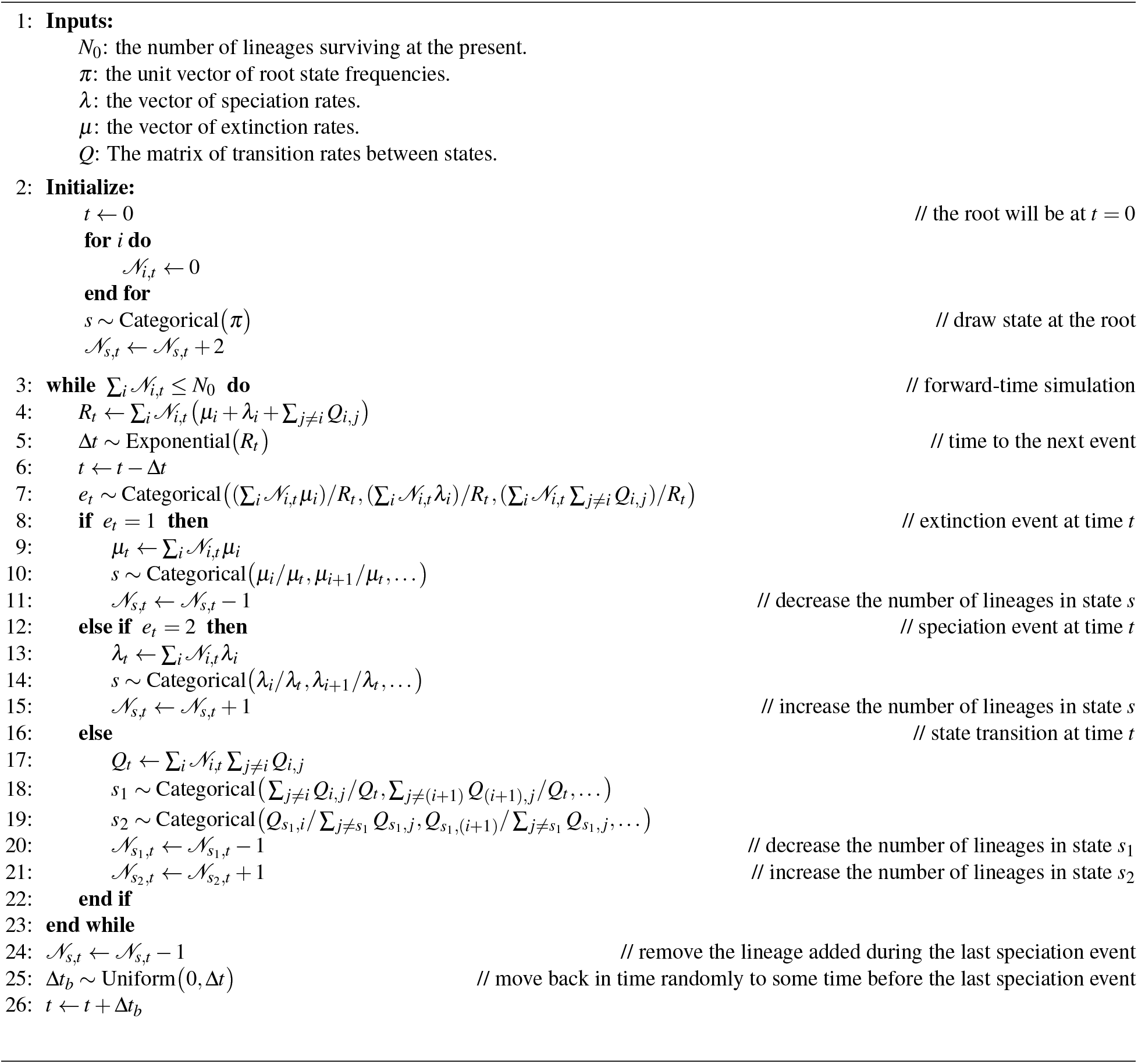

Another option would be to condition on the number of tips with each character state value, 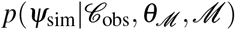, where 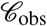 is the observed character data. This will generate a more constrained set of trees than conditioning only on the total number of tips, without regard for their states. The procedure for PPS conditioned on number of tips in each state is based on Bromham et al. (2016) and described in Algorithm 3.

#### Algorithm 3 Backward-time simulation of the state-dependent birth-death process conditioned on the number of lineages in each state at the present. 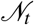 is the vector of the number of lineages in each state at time *t*. The root age is at time *t_r_* > ^0^ and the present is at time *t* = 0.

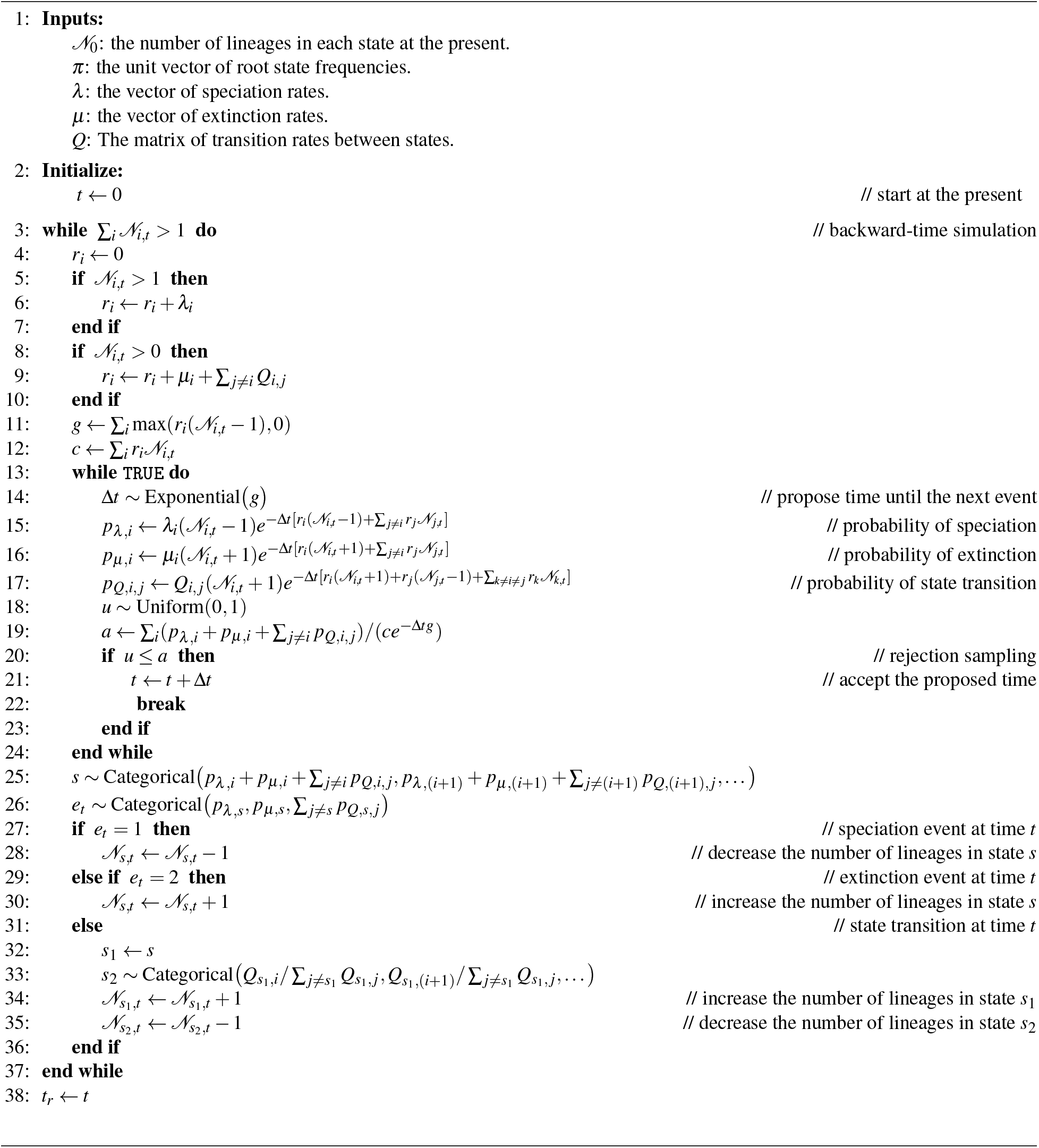

Finally, we could constrain the distribution of simulated data even more by conditioning on the entire observed tree and tip states. This is essentially stochastic mapping under a model of trait-dependent diversification, as described in Freyman and Höhna (2019).

**Figure S1:**
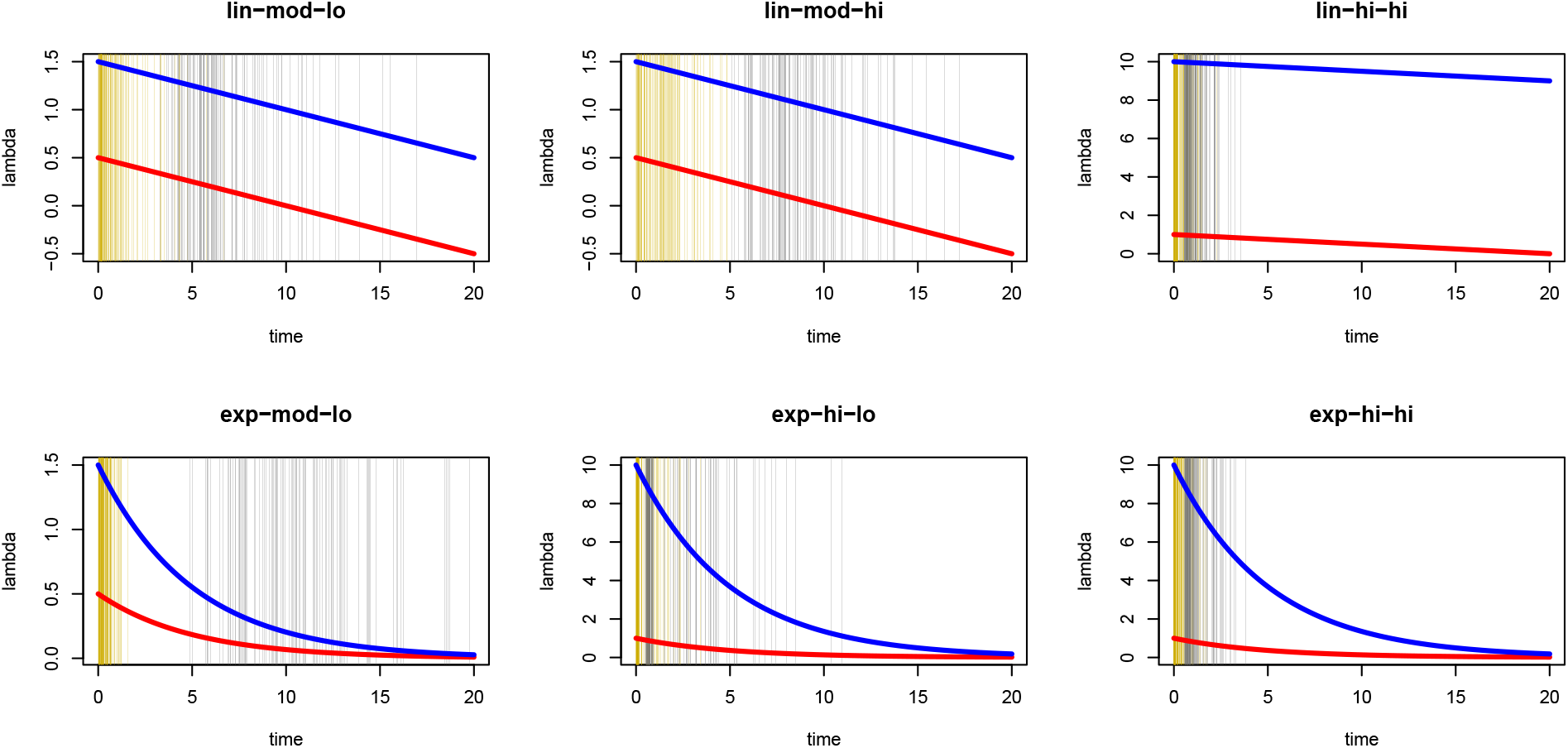
Rates through time for tdBiSSE scenarios. Red and blue lines are speciation rates of lineages in state 0 and 1 respectively; golden and gray vertical lines are root and tip times respectively of the 100 simulated trees.

**Table S1:** All summary statistics used. The ‘Variant,’ ‘Delta,’ and ‘Conditioning’ columns are explained in Table 1. When explicit names of each statistic are used elsewhere, they are composed of the first four columns here, `e.g., branch_length_max_absdelta_ntips` or Colless_index_ntips. Statistics ‘Retained’ were the ones which consistently inferred the CID-1 and BiSSE models to be adequate for trees generated under the same respective model. The different aspects of the phylogeny—topology, branch lengths (node ages), trait states at the tips, and reconstruction of ancestral trait states—that contribute to each statistic are indicated by the remaining columns. (Table continues across multiple pages.)

| Statistic | Variant | Delta | Conditioning | Retained | Topology | BrLens | TipSts | StRecon |
| --- | --- | --- | --- | --- | --- | --- | --- | --- |
| branch length | max | absdelta | ntips |  |  | x | x |  |
| branch length | max | absdelta | states | x |  | x | x |  |
| branch length | max | absdelta | time |  |  | x | x |  |
| branch length | max | absdelta | tree |  |  | x | x |  |
| branch length | max | delta | ntips |  |  | x | x |  |
| branch length | max | delta | states |  |  | x | x |  |
| branch length | max | delta | time |  |  | x | x |  |
| branch length | max | delta | tree |  |  | x | x |  |
| branch length | median | absdelta | ntips |  |  | x | x |  |
| branch length | median | absdelta | states |  |  | x | x |  |
| branch length | median | absdelta | time |  |  | x | x |  |
| branch length | median | absdelta | tree |  |  | x | x |  |
| branch length | median | delta | ntips |  |  | x | x |  |
| branch length | median | delta | states |  |  | x | x |  |
| branch length | median | delta | time |  |  | x | x |  |
| branch length | median | delta | tree | x |  | x | x |  |
| branch length | min | absdelta | ntips | x |  | x | x |  |
| branch length | min | absdelta | states |  |  | x | x |  |
| branch length | min | absdelta | time | x |  | x | x |  |
| branch length | min | absdelta | tree |  |  | x | x |  |
| branch length | min | delta | ntips |  |  | x | x |  |
| branch length | min | delta | states |  |  | x | x |  |
| branch length | min | delta | time | x |  | x | x |  |
| branch length | min | delta | tree |  |  | x | x |  |
| branch length | var | absdelta | ntips |  |  | x | x |  |
| branch length | var | absdelta | states |  |  | x | x |  |
| branch length | var | absdelta | time |  |  | x | x |  |
| branch length | var | absdelta | tree |  |  | x | x |  |
| branch length | var | delta | ntips | x |  | x | x |  |
| branch length | var | delta | states |  |  | x | x |  |
| branch length | var | delta | time | x |  | x | x |  |
| branch length | var | delta | tree |  |  | x | x |  |
| Colless index |  | absdelta | ntips |  | x |  | x |  |
| Colless index |  | absdelta | states |  | x |  | x |  |
| Colless index |  | absdelta | time |  | x |  | x |  |
| Colless index |  | absdelta | tree |  | x |  | x |  |
| Colless index |  | delta | ntips | x | x |  | x |  |
| Colless index |  | delta | states |  | x |  | x |  |
| Colless index |  | delta | time | x | x |  | x |  |
| Colless index |  | delta | tree | x | x |  | x |  |
| Colless index |  |  | ntips |  | x |  |  |  |
| Colless index |  |  | states |  | x |  |  |  |
| Colless index |  |  | time |  | x |  |  |  |
| Colless index |  |  | tree |  | x |  |  |  |
| EDR | max | absdelta | ntips |  |  | x | x |  |
| EDR | max | absdelta | states | x |  | x | x |  |
| EDR | max | absdelta | time |  |  | x | x |  |
| EDR | max | absdelta | tree | x |  | x | x |  |
| EDR | max | delta | ntips |  |  | x | x |  |
| EDR | max | delta | states |  |  | x | x |  |
| EDR | max | delta | time | x |  | x | x |  |
| EDR | max | delta | tree | x |  | x | x |  |
| EDR | median | absdelta | ntips |  |  | x | x |  |
| EDR | median | absdelta | states |  |  | x | x |  |
| EDR | median | absdelta | time |  |  | x | x |  |
| EDR | median | absdelta | tree |  |  | x | x |  |
| EDR | median | delta | ntips | x |  | x | x |  |
| EDR | median | delta | states |  |  | x | x |  |
| EDR | median | delta | time |  |  | x | x |  |
| EDR | median | delta | tree | x |  | x | x |  |
| EDR | min | absdelta | ntips |  |  | x | x |  |
| EDR | min | absdelta | states |  |  | x | x |  |
| EDR | min | absdelta | time | x |  | x | x |  |
| EDR | min | absdelta | tree |  |  | x | x |  |
| EDR | min | delta | ntips |  |  | x | x |  |
| EDR | min | delta | states |  |  | x | x |  |
| EDR | min | delta | time |  |  | x | x |  |
| EDR | min | delta | tree |  |  | x | x |  |
| EDR | var | absdelta | ntips |  |  | x | x |  |
| EDR | var | absdelta | states |  |  | x | x |  |
| EDR | var | absdelta | time |  |  | x | x |  |
| EDR | var | absdelta | tree |  |  | x | x |  |
| EDR | var | delta | ntips | x |  | x | x |  |
| EDR | var | delta | states |  |  | x | x |  |
| EDR | var | delta | time | x |  | x | x |  |
| EDR | var | delta | tree | x |  | x | x |  |
| FiSSE |  |  | ntips | x | x | x | x |  |
| FiSSE |  |  | states |  | x | x | x |  |
| FiSSE |  |  | time | x | x | x | x |  |
| FiSSE |  |  | tree | x | x | x | x |  |
| Fitch score |  |  | ntips |  | x |  | x | x |
| Fitch score |  |  | states |  | x |  | x | x |
| Fitch score |  |  | time |  | x |  | x | x |
| Fitch score |  |  | tree |  | x |  | x | x |
| gamma statistic |  | absdelta | ntips |  |  | x | x |  |
| gamma statistic |  | absdelta | states |  |  | x | x |  |
| gamma statistic |  | absdelta | time |  |  | x | x |  |
| gamma statistic |  | absdelta | tree |  |  | x | x |  |
| gamma statistic |  | delta | ntips |  |  | x | x |  |
| gamma statistic |  | delta | states |  |  | x | x |  |
| gamma statistic |  | delta | time |  |  | x | x |  |

Table S1: (continued)
| Statistic | Variant | Delta | Conditioning | Retained | Topology | BrLens | TipSts | StRecon |
| --- | --- | --- | --- | --- | --- | --- | --- | --- |
| gamma statistic |  | delta | tree |  |  | x | x |  |
| gamma statistic |  |  | ntips | x |  | x |  |  |
| gamma statistic |  |  | states |  |  | x |  |  |
| gamma statistic |  |  | time | x |  | x |  |  |
| gamma statistic |  |  | tree |  |  | x |  |  |
| InverseES | max | absdelta | ntips |  | x | x | x |  |
| InverseES | max | absdelta | states |  | x | x | x |  |
| InverseES | max | absdelta | time |  | x | x | x |  |
| InverseES | max | absdelta | tree |  | x | x | x |  |
| InverseES | max | delta | ntips |  | x | x | x |  |
| InverseES | max | delta | states | x | x | x | x |  |
| InverseES | max | delta | time | x | x | x | x |  |
| InverseES | max | delta | tree | x | x | x | x |  |
| InverseES | median | absdelta | ntips |  | x | x | x |  |
| InverseES | median | absdelta | states |  | x | x | x |  |
| InverseES | median | absdelta | time |  | x | x | x |  |
| InverseES | median | absdelta | tree |  | x | x | x |  |
| InverseES | median | delta | ntips | x | x | x | x |  |
| InverseES | median | delta | states |  | x | x | x |  |
| InverseES | median | delta | time | x | x | x | x |  |
| InverseES | median | delta | tree |  | x | x | x |  |
| InverseES | min | absdelta | ntips |  | x | x | x |  |
| InverseES | min | absdelta | states |  | x | x | x |  |
| InverseES | min | absdelta | time |  | x | x | x |  |
| InverseES | min | absdelta | tree |  | x | x | x |  |
| InverseES | min | delta | ntips |  | x | x | x |  |
| InverseES | min | delta | states |  | x | x | x |  |
| InverseES | min | delta | time |  | x | x | x |  |
| InverseES | min | delta | tree |  | x | x | x |  |
| InverseES | var | absdelta | ntips |  | x | x | x |  |
| InverseES | var | absdelta | states |  | x | x | x |  |
| InverseES | var | absdelta | time |  | x | x | x |  |
| InverseES | var | absdelta | tree |  | x | x | x |  |
| InverseES | var | delta | ntips | x | x | x | x |  |
| InverseES | var | delta | states | x | x | x | x |  |
| InverseES | var | delta | time | x | x | x | x |  |
| InverseES | var | delta | tree |  | x | x | x |  |
| MNTD |  | absdelta | ntips |  | x | x | x |  |
| MNTD |  | absdelta | states |  | x | x | x |  |
| MNTD |  | absdelta | time |  | x | x | x |  |
| MNTD |  | absdelta | tree |  | x | x | x |  |
| MNTD |  | delta | ntips | x | x | x | x |  |
| MNTD |  | delta | states |  | x | x | x |  |
| MNTD |  | delta | time | x | x | x | x |  |
| MNTD |  | delta | tree |  | x | x | x |  |
| MNTD | mean |  | ntips | x | x | x | x |  |
| MNTD | mean |  | states |  | x | x | x |  |
| MNTD | mean |  | time | x | x | x | x |  |
| MNTD | mean |  | tree |  | x | x | x |  |
| MNTD node |  | absdelta | ntips |  | x | x | x |  |
| MNTD node |  | absdelta | states |  | x | x | x |  |

Table S1: (continued)
| Statistic | Variant | Delta | Conditioning | Retained | Topology | BrLens | TipSts | StRecon |
| --- | --- | --- | --- | --- | --- | --- | --- | --- |
| MNTD node |  | absdelta | time |  | x | x | x |  |
| MNTD node |  | absdelta | tree |  | x | x | x |  |
| MNTD node |  | delta | ntips | x | x | x | x |  |
| MNTD node |  | delta | states |  | x | x | x |  |
| MNTD node |  | delta | time | x | x | x | x |  |
| MNTD node |  | delta | tree | x | x | x | x |  |
| MNTD node | mean |  | ntips | x | x | x | x |  |
| MNTD node | mean |  | states |  | x | x | x |  |
| MNTD node | mean |  | time | x | x | x | x |  |
| MNTD node | mean |  | tree |  | x | x | x |  |
| MPD |  | absdelta | ntips | x | x | x | x |  |
| MPD |  | absdelta | states |  | x | x | x |  |
| MPD |  | absdelta | time | x | x | x | x |  |
| MPD |  | absdelta | tree |  | x | x | x |  |
| MPD |  | delta | ntips |  | x | x | x |  |
| MPD |  | delta | states | x | x | x | x |  |
| MPD |  | delta | time |  | x | x | x |  |
| MPD |  | delta | tree | x | x | x | x |  |
| MPD | mean |  | ntips | x | x | x | x |  |
| MPD | mean |  | states |  | x | x | x |  |
| MPD | mean |  | time | x | x | x | x |  |
| MPD | mean |  | tree | x | x | x | x |  |
| MPD node |  | absdelta | ntips |  | x | x | x |  |
| MPD node |  | absdelta | states |  | x | x | x |  |
| MPD node |  | absdelta | time |  | x | x | x |  |
| MPD node |  | absdelta | tree | x | x | x | x |  |
| MPD node |  | delta | ntips |  | x | x | x |  |
| MPD node |  | delta | states | x | x | x | x |  |
| MPD node |  | delta | time | x | x | x | x |  |
| MPD node |  | delta | tree | x | x | x | x |  |
| MPD node | mean |  | ntips | x | x | x | x |  |
| MPD node | mean |  | states | x | x | x | x |  |
| MPD node | mean |  | time | x | x | x | x |  |
| MPD node | mean |  | tree |  | x | x | x |  |
| node age | max | absdelta | ntips |  |  | x | x |  |
| node age | max | absdelta | states |  |  | x | x |  |
| node age | max | absdelta | time |  |  | x | x |  |
| node age | max | absdelta | tree |  |  | x | x |  |
| node age | max | delta | ntips |  |  | x | x |  |
| node age | max | delta | states |  |  | x | x |  |
| node age | max | delta | time |  |  | x | x |  |
| node age | max | delta | tree |  |  | x | x |  |
| node age | median | absdelta | ntips | x |  | x | x |  |
| node age | median | absdelta | states |  |  | x | x |  |
| node age | median | absdelta | time | x |  | x | x |  |
| node age | median | absdelta | tree | x |  | x | x |  |
| node age | median | delta | ntips |  |  | x | x |  |
| node age | median | delta | states |  |  | x | x |  |
| node age | median | delta | time | x |  | x | x |  |
| node age | median | delta | tree | x |  | x | x |  |

Table S1: (continued)
| Statistic | Variant | Delta | Conditioning | Retained | Topology | BrLens | TipSts | StRecon |
| --- | --- | --- | --- | --- | --- | --- | --- | --- |
| node age | min | absdelta | ntips |  |  | x | x |  |
| node age | min | absdelta | states |  |  | x | x |  |
| node age | min | absdelta | time |  |  | x | x |  |
| node age | min | absdelta | tree | x |  | x | x |  |
| node age | min | delta | ntips |  |  | x | x |  |
| node age | min | delta | states |  |  | x | x |  |
| node age | min | delta | time | x |  | x | x |  |
| node age | min | delta | tree |  |  | x | x |  |
| node age | var | absdelta | ntips |  |  | x | x |  |
| node age | var | absdelta | states |  |  | x | x |  |
| node age | var | absdelta | time |  |  | x | x |  |
| node age | var | absdelta | tree |  |  | x | x |  |
| node age | var | delta | ntips | x |  | x | x |  |
| node age | var | delta | states |  |  | x | x |  |
| node age | var | delta | time | x |  | x | x |  |
| node age | var | delta | tree |  |  | x | x |  |
| num lineages surviving |  |  | time | x |  |  |  |  |
| PSSP | max | absdelta | ntips |  | x | x | x | x |
| PSSP | max | absdelta | states |  | x | x | x | x |
| PSSP | max | absdelta | time |  | x | x | x | x |
| PSSP | max | absdelta | tree |  | x | x | x | x |
| PSSP | max | delta | ntips |  | x | x | x | x |
| PSSP | max | delta | states | x | x | x | x | x |
| PSSP | max | delta | time |  | x | x | x | x |
| PSSP | max | delta | tree |  | x | x | x | x |
| PSSP | median | absdelta | ntips | x | x | x | x | x |
| PSSP | median | absdelta | states |  | x | x | x | x |
| PSSP | median | absdelta | time |  | x | x | x | x |
| PSSP | median | absdelta | tree |  | x | x | x | x |
| PSSP | median | delta | ntips | x | x | x | x | x |
| PSSP | median | delta | states |  | x | x | x | x |
| PSSP | median | delta | time | x | x | x | x | x |
| PSSP | median | delta | tree | x | x | x | x | x |
| PSSP | min | absdelta | ntips | x | x | x | x | x |
| PSSP | min | absdelta | states |  | x | x | x | x |
| PSSP | min | absdelta | time | x | x | x | x | x |
| PSSP | min | absdelta | tree |  | x | x | x | x |
| PSSP | min | delta | ntips |  | x | x | x | x |
| PSSP | min | delta | states |  | x | x | x | x |
| PSSP | min | delta | time | x | x | x | x | x |
| PSSP | min | delta | tree |  | x | x | x | x |
| PSSP | sum | absdelta | ntips |  | x | x | x | x |
| PSSP | sum | absdelta | states |  | x | x | x | x |
| PSSP | sum | absdelta | time |  | x | x | x | x |
| PSSP | sum | absdelta | tree |  | x | x | x | x |
| PSSP | sum | delta | ntips | x | x | x | x | x |
| PSSP | sum | delta | states |  | x | x | x | x |
| PSSP | sum | delta | time | x | x | x | x | x |
| PSSP | sum | delta | tree | x | x | x | x | x |
| PSSP | var | absdelta | ntips | x | x | x | x | x |
| PSSP | var | absdelta | states |  | x | x | x | x |

Table S1: (continued)
| Statistic | Variant | Delta | Conditioning | Retained | Topology | BrLens | TipSts | StRecon |
| --- | --- | --- | --- | --- | --- | --- | --- | --- |
| PSSP | var | absdelta | time |  | x | x | x | x |
| PSSP | var | absdelta | tree |  | x | x | x | x |
| PSSP | var | delta | ntips |  | x | x | x | x |
| PSSP | var | delta | states |  | x | x | x | x |
| PSSP | var | delta | time | x | x | x | x | x |
| PSSP | var | delta | tree | x | x | x | x | x |
| state relative time |  | absdelta | ntips |  | x | x | x | x |
| state relative time |  | absdelta | states |  | x | x | x | x |
| state relative time |  | absdelta | time |  | x | x | x | x |
| state relative time |  | absdelta | tree |  | x | x | x | x |
| state relative time |  | delta | ntips | x | x | x | x | x |
| state relative time |  | delta | states |  | x | x | x | x |
| state relative time |  | delta | time | x | x | x | x | x |
| state relative time |  | delta | tree | x | x | x | x | x |
| tip state frequencies |  | absdelta | ntips |  |  |  | x |  |
| tip state frequencies |  | absdelta | tree |  |  |  | x |  |
| tip state frequencies |  | delta | ntips | x |  |  | x |  |
| tip state frequencies |  | delta | tree | x |  |  | x |  |
| tree length |  | absdelta | ntips |  |  | x | x |  |
| tree length |  | absdelta | states |  |  | x | x |  |
| tree length |  | absdelta | time | x |  | x | x |  |
| tree length |  | absdelta | tree |  |  | x | x |  |
| tree length |  | delta | ntips | x |  | x | x |  |
| tree length |  | delta | states |  |  | x | x |  |
| tree length |  | delta | time | x |  | x | x |  |
| tree length |  | delta | tree | x |  | x | x |  |

## Notes

### Competing Interest Statement

The authors have declared no competing interest.

